# GENETIC AND PHARMACOLOGIC ACTIVATION OF BECLIN1 PREVENTS ALDOSTERONE-INDUCED CARDIOVASCULAR DAMAGE

**DOI:** 10.64898/2026.07.22.740044

**Authors:** Rafael M Costa, Ariane Bruder, Juliano V Alves, Debora M Cerqueira, Luis Oliveira de Moraes, Tyler Beling, Sergio Guerrero, Jaqueline Ho, Rita de Cassia A Tostes, Thiago Bruder-Nascimento

## Abstract

**Background:** Aldosterone promotes endothelial dysfunction and cardiovascular injury through mineralocorticoid receptor (MR) activation. Autophagy is essential for endothelial homeostasis, yet its role in aldosterone-mediated vascular dysfunction remains unclear. We tested whether aldosterone impairs autophagic flux and whether restoring autophagy via Beclin1 (BCN1) activation protects vascular and cardiac function.

**Methods:** Endothelial and vascular responses to aldosterone were assessed in wild-type mice, BCN1 gain-of-function mice (Becn1), and mice treated with spermidine or a BCN1- activating TB-peptide. Vascular function, nitric oxide (NO)/reactive oxygen species (ROS) production, autophagy markers, endothelial migration, and cardiac fibrosis were evaluated using wire myography, fluorescence assays, Western blotting, confocal microscopy, migration assays, and histology.

**Results:** Aldosterone impaired endothelium-dependent relaxation, decreased NO, increased ROS, and disrupted autophagic flux in an MR-dependent manner, indicated by LC3 accumulation and reduced p62 and BCN1 expression. Spermidine restored endothelial function and normalized NO and ROS levels. BCN1 gain-of-function mice were protected from aldosterone-induced endothelial dysfunction and exhibited reduced coronary and myocardial fibrosis. TB-peptide activation of BCN1 enhanced autophagic flux, improved vascular function, decreased cardiac fibrosis, and rescued endothelial migration impaired by aldosterone.

**Conclusions:** Aldosterone induces endothelial dysfunction by suppressing autophagic flux through MR activation. Genetic or pharmacologic enhancement of BCN1-dependent autophagy restores endothelial homeostasis and prevents vascular and cardiac injury, identifying autophagy activation as a promising therapeutic approach for cardiovascular diseases associated with mineralocorticoid excess.

## Introduction

Aldosterone, acting through the mineralocorticoid receptor (MR), plays a central role in the development of vascular dysfunction and cardiovascular disease. Excess aldosterone promotes endothelial oxidative stress, reduces nitric oxide (NO) bioavailability, and induces inflammation and fibrosis, contributing to impaired vasomotor function and end-organ injury^1–3^. Although MR antagonists provide clinical benefit^4, 5^, the cellular mechanisms linking aldosterone signaling to endothelial dysfunction and injury remain incompletely understood. Emerging evidence indicates that endothelial autophagy, a fundamental homeostatic process^5^, is disrupted by aldosterone exposure^6^, yet the functional consequences of this defect and its therapeutic relevance have not been fully elucidated.

Autophagy is a fundamental homeostatic mechanism that preserves endothelial function by maintaining mitochondrial quality control, redox balance, and cellular repair capacity^7–10^. Its dysregulation contributes to vascular stiffening, impaired angiogenesis, and microvascular rarefaction during cardiovascular stress^11, 12^. Among the core autophagy regulators, Beclin1 (BCN1) functions as a scaffold for autophagosome initiation and is critical for maintaining autophagy flux in endothelial cells. Alterations in BCN1-dependent autophagy have been implicated in aging, metabolic dysfunction, and vascular injury^9, 13, 14^, suggesting that restoring autophagy may have therapeutic potential in aldosterone-driven disease.

Pharmacologic activation of autophagy has emerged as a promising approach to improving endothelial health. Spermidine, an endogenous polyamine and autophagy inducer, has been shown to promote cardiovascular resilience, prolong lifespan, and enhance endothelial NO signaling^9, 15, 16^. However, whether spermidine can counteract aldosterone-induced endothelial dysfunction and the associated disturbances in redox homeostasis remains to be determined. Likewise, targeted enhancement of autophagy through direct BCN1 activation represents a novel therapeutic strategy. A recently developed BCN1-activating peptide (TB-peptide) amplifies autophagy flux and improves cellular stress resistance, offering a mechanistically specific tool to restore impaired autophagy in disease settings^9, 17, 18^.

Beyond pharmacological activation, genetic augmentation of autophagy provides an opportunity to define the mechanistic importance of BCN1 *in vivo*. Mice engineered to overexpress BCN1 exhibit enhanced autophagy flux, improved metabolic homeostasis, and resistance to cardiac stressors and perivascular adipose (PVAT) tissue dysfunction, and partially protection against aldosterone-induced hypertension^9, 18^. Whether genetic BCN1 overactivation protects against aldosterone-induced vascular dysfunction in small vessels and cardiac injury has not been explored and could help clarify the causal role of autophagy in this context.

Given the central role of endothelial cells in regulating vascular tone, repair, and redox balance, elucidating how BCN1-dependent autophagy modulates endothelial responses to aldosterone is of particular interest. Preservation of NO bioavailability and suppression of oxidative stress are critical features of endothelial resilience, while endothelial migration and regenerative capacity are essential for maintaining vascular integrity following injury. Understanding how BCN1 activation influences these functions may reveal a unifying mechanism linking autophagy restoration to protection against aldosterone-induced vascular and cardiac pathology.

We hypothesized that aldosterone impairs endothelial autophagic flux through mineralocorticoid receptor activation and that restoring BCN1-dependent autophagy, either pharmacologically or genetically, protects against vascular dysfunction and cardiac injury. Our findings demonstrate that aldosterone disrupts endothelial autophagy flux in an MR-dependent manner, leading to increased reactive oxygen species (ROS), reduced NO availability, diminished endothelial migration, and vascular dysfunction. Restoring autophagy through spermidine treatment or BCN1 activation preserved endothelial function and prevented both vascular dysfunction and cardiac fibrosis. These results identify BCN1-dependent autophagy as a central protective mechanism against aldosterone-induced cardiovascular injury and highlight autophagy activation as a potential therapeutic strategy.

## Material and Methods

### Animals

Male wild-type (WT; C57BL/6) mice and BCN1 gain-of-function mice [*Becn1*^F121A^ knock-in mice (B6.129(Cg)-*Becn1*^tm2.1Blev^/J, Jackson Laboratory, Stock No. 033360] aged ^10–12^ weeks were used for all experiments. Animals were maintained on standard chow with ad libitum access to tap water. All procedures were approved by the Institutional Animal Care and Use Committee (IACUC) at the University of South Alabama (Protocol 2219557) and the University of Pittsburgh (Protocols 19065333 and 22061179), and conformed to the Guide for the Care and Use of Laboratory Animals. Experiments were conducted in the Department of Pediatrics at the University of Pittsburgh and the Department of Physiology and Cell Biology at the University of South Alabama. Mice were euthanized by gradual-fill CO₂ inhalation (20– 30% chamber volume per minute) followed by cervical dislocation as a secondary method to ensure death.

### Aldosterone Infusion *In Vivo*

Mice were infused for 14 days with either vehicle (saline) or aldosterone (600 μg/kg/day; Sigma-Aldrich, A9477) delivered via subcutaneously implanted ALZET osmotic mini-pumps (Model 2002). Throughout the infusion period, all animals received 1% NaCl in their drinking water, as previously described ^2, 3^.

For surgical procedures, mice received extended-release buprenorphine (Ethiqa XR, 3.25 mg/kg, subcutaneous) for analgesia. Ethiqa XR is supplied at a concentration of 1.3 mg/mL, corresponding to a dosing volume of 2.5 mL/kg (e.g., approximately 0.063 mL for a 25 g mouse).

### Spermidine Treatment *In Vivo*

To evaluate the effects of autophagy induction, a subset of mice received spermidine (3 mM in drinking water; Sigma-Aldrich), as described previously^15^, beginning on day 7 of aldosterone infusion and continuing for 7 days.

### TB-Peptide Treatment *In Vivo*

Pharmacologic activation of BCN1 was achieved using TB-peptide (Sigma). After 7 days of aldosterone infusion, mice received TB-peptide (16 mg/kg/day, intraperitoneally) for an additional 7 days^9^.

### Vascular Reactivity in Mesenteric Arteries

Vascular function was assessed in second-order mesenteric arteries following established protocols^3^, ^19^. Vessels were dissected free of connective tissue, cut into 2-mm rings, and mounted on a wire myograph (Danish Myo Technology) for isometric tension recording (PowerLab, AD Instruments). Rings were bathed in Krebs–Henseleit solution (37°C, aerated with 95% O₂/5% CO₂; composition in mM: 130 NaCl, 4.7 KCl, 1.17 MgSO₄, 0.03 EDTA, 1.6 CaCl₂, 14.9 NaHCO₃, 1.18 KH₂PO₄, 5.5 glucose). After a 30-min equilibration period, vascular viability was confirmed with 60 mM KCl. Concentration–response curves were generated for phenylephrine (PE): α₁-adrenergic vasoconstriction, acetylcholine (ACh): endothelium-dependent relaxation, and sodium nitroprusside (SNP): endothelium-independent relaxation.

### Cardiac Histology

Hearts were harvested and fixed in 4% paraformaldehyde for 12 hours, then transferred to 70% ethanol. Samples were paraffin-embedded, sectioned, and stained with Masson’s trichrome for analysis of cardiac fibrosis.

### Mouse Endothelial Cell Culture and Treatments

Mouse mesenteric endothelial cells (MECs) (Cell Biologics #C57-6055) were cultured in Complete Mouse Endothelial Cell Medium supplemented with the manufacturer’s supplement kit. Cells between passages ^4–8^ were used. MECs were exposed to aldosterone (0.1 μM, 3 h). For autophagy activation experiments, cells were pre-incubated with spermidine (3 nM) or TB-peptide (30 μM) prior to aldosterone treatment.

### Immunofluorescence

MECs were fixed in 4% paraformaldehyde for 1 hour and subsequently rinsed three times with PBS for 10 minutes each. Cells were then blocked in 3% BSA for 1 hour and incubated overnight at 4°C with an anti-LC3 primary antibody (Catalog number, Company). The following day, samples were washed with PBS and incubated for 2 hours with an Alexa Fluor 594–conjugated anti-rabbit secondary antibody (#711-587-003, Jackson ImmunoResearch Laboratories, West Grove, PA, USA). After additional PBS washes, slides were mounted using Fluoro Gel with DABCO™ (Electron Microscopy Science, Hatfield, PA, USA). All antibodies were used at a 1:100 dilution. Fluorescent images were acquired using a Leica Stellaris 5 confocal microscope equipped with silicon HyD detectors (Leica, Buffalo Grove, IL, USA). Imaging was performed at the Cell Imaging Core Laboratory in the John G. Rangos Research Center, UPMC Children’s Hospital (RRID:SCR_025132).

### Western Blot Analysis

Protein was isolated from mesenteric arteries using RIPA buffer (30 mM HEPES, pH 7.4; 150 mM NaCl; 1% NP-40; 0.5% sodium deoxycholate; 0.1% SDS; 5 mM EDTA) supplemented with protease and phosphatase inhibitors. Lysates were centrifuged at 15,000 rpm for 10 min, and supernatants collected. For aortic tissue, 25 μg protein was used. MECs were lysed directly in 2× Laemmli Buffer with β-mercaptoethanol. Proteins were resolved on polyacrylamide gradient gels (Bio-Rad), transferred to PVDF membranes, and blocked in 5% milk or 1% BSA. Membranes were incubated overnight at 4°C with primary antibodies (Supplementary Table 1), followed by HRP-conjugated secondary antibodies. Bands were visualized using enhanced chemiluminescence (SuperSignal West Femto, Thermo Scientific).

### Nitric Oxide Quantification

NO production was assessed using the DAF-2 DA fluorescent probe (5 μM; Thermo Scientific). MECs were treated with aldosterone (0.1 μM, 24 h) with or without TB-peptide (30 μM). After staining with DAF-2 DA for 30 min, fluorescence was measured using a SpectraMax i3x reader (Ex 485 nm/Em 538 nm).

### Reactive Oxygen Species Measurements

#### Amplex Red Assay (H_2_O_2_ Production)

MECs treated with aldosterone with or without TB-peptide were lysed and centrifuged at 1,000 g for 5 min at 4°C. Aortic tissue lysates were incubated with Amplex Red reagent (0.1 mM) and horseradish peroxidase (0.35 U/mL) in buffer (25 mM HEPES, pH 7.4; 120 mM NaCl; 3 mM KCl; 1 mM MgCl₂) with or without catalase (300 U/mL). Fluorescence was detected using a Synergy 4 reader (Ex 530/25 nm; Em 590/35 nm) and normalized to protein content (Bradford assay).

#### Lucigenin Chemiluminescence (Superoxide Production)

MECs were harvested in lysis buffer (20 mM KH₂PO₄, 1 mM EGTA, protease inhibitors). Samples were mixed with assay buffer (50 mM KH₂PO₄, 1 mM EGTA, 150 mM sucrose) containing lucigenin (5 μM). Basal luminescence was recorded, followed by stimulation with NADPH (10⁻ M). Luminescence was measured for 30 cycles of 30 s each. Data were normalized to protein content and expressed as % of control relative light units (RLU).

### Endothelial Cell Migration Assays

#### Transwell Migration

We used transwell inserts (8-μm pore size; Costar) as described before^20^. MECs (2.5×10^4^ cells/well) were plated in the upper chamber in medium containing 0.5% FBS. The lower chamber contained aldosterone (0.1 μM) with or without TB-peptide (30 μM). After 12 h, migrated cells were fixed with 4% formaldehyde, stained with trypan blue, and imaged using an Echo Revolve microscope.

### Scratch Wound Assay

MECs (1 × 10 cells/mL) were plated in 12-well plates and grown overnight. After 2 h of serum starvation (0.5% FBS), a linear scratch was made using a 200-μL pipette tip. Detached cells were removed with PBS. Fresh medium containing aldosterone (0.1 μM) with or without TB-peptide (30 μM) was added, and migration was monitored for 24 h. Images were taken with an Echo Revolve microscope and quantified using ImageJ.

### Statistical Analysis

Data are expressed as mean ± SEM. Comparisons among three groups were performed using one-way ANOVA followed by Tukey’s post hoc test. When four groups or two experimental factors were analyzed, two-way ANOVA with Tukey’s post hoc test was used.

Differences between two groups were assessed using an unpaired Student’s t-test. Concentration–response curves were analyzed by nonlinear regression to calculate maximal responses (E_max) and pD₂ values. All statistical analyses were performed using GraphPad Prism 10 (GraphPad Software, San Diego, CA). A value of p < 0.05 was considered statistically significant.

## Results

### Aldosterone causes vascular dysfunction and disrupts endothelial autophagy flux in a MR-dependent response

We first assessed whether aldosterone-induced vascular dysfunction is associated with alterations in autophagy signaling. In mesenteric arteries, aldosterone treatment induced marked vascular dysfunction, characterized by enhanced PE-induced contractility and impaired ACh-mediated endothelium-dependent relaxation in mesenteric arteries (Figure 1A– B). In MECs, aldosterone exposure reduced NO production and increased ROS generation (Figure 1C-E). Notably, these functional and redox disturbances were accompanied by disrupted autophagic flux, reflected by LC3I/II accumulation (Figure 1F-G) and decreased expression of p62 and BCN1 (Figure 1H-I, respectively).

**Figure 1.**
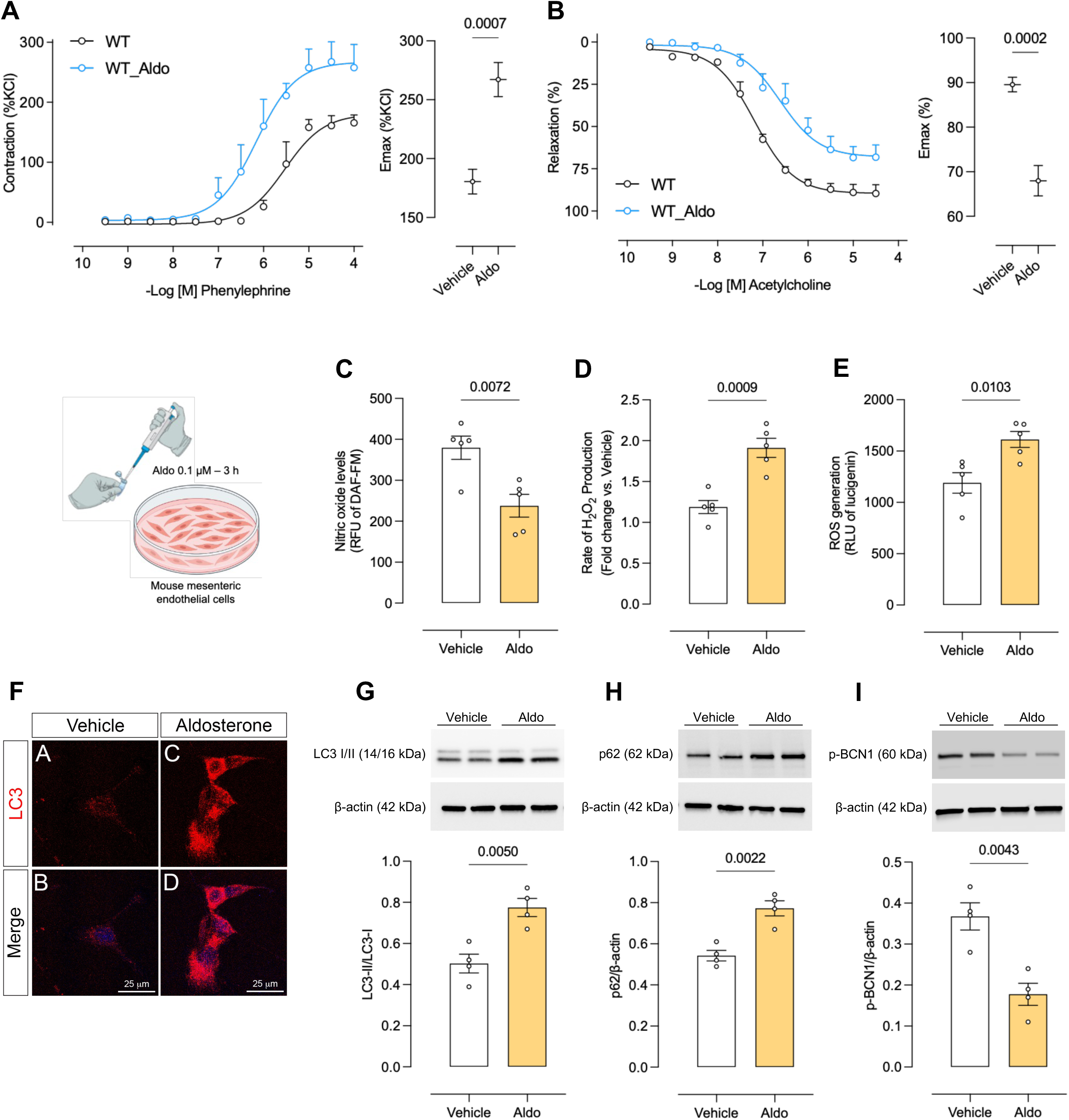
Aldosterone impairs endothelial function and modulates autophagy-related signaling in endothelial cells. Concentration–response curves to phenylephrine and acetylcholine in second-order mesenteric resistance arteries from male (A and B) wild-type mice treated with vehicle or aldosterone (600 µg/kg/day for 14 days). Nitric oxide production (C), hydrogen peroxide levels (D), and reactive oxygen species generation (E) in mesenteric endothelial cells (MEC) treated with vehicle or aldosterone (0.1 µM, 3 h). Representative immunofluorescence images of LC3 (red) and nuclei (DAPI, blue) in MEC treated with vehicle or aldosterone (F). Representative Western blot images and densitometric quantification of LC3 I/II (G), p62 (H), and phosphorylated Beclin-1 (p-BCN1) (I) expression in MEC treated with vehicle or aldosterone. Values represent mean ± SEM (n = 4–7). Statistical analysis was performed using ANOVA.

To determine whether aldosterone-induced autophagic flux impairment occurs through MR activation, MECs were treated with aldosterone in the presence of the MR antagonist eplerenone. Eplerenone completely prevented the aldosterone-induced alterations in LC3I/II, p62, and BCN1 expression, indicating that aldosterone disrupts autophagy signaling in an MR-dependent manner (Figure 2A–C).

**Figure 2.**
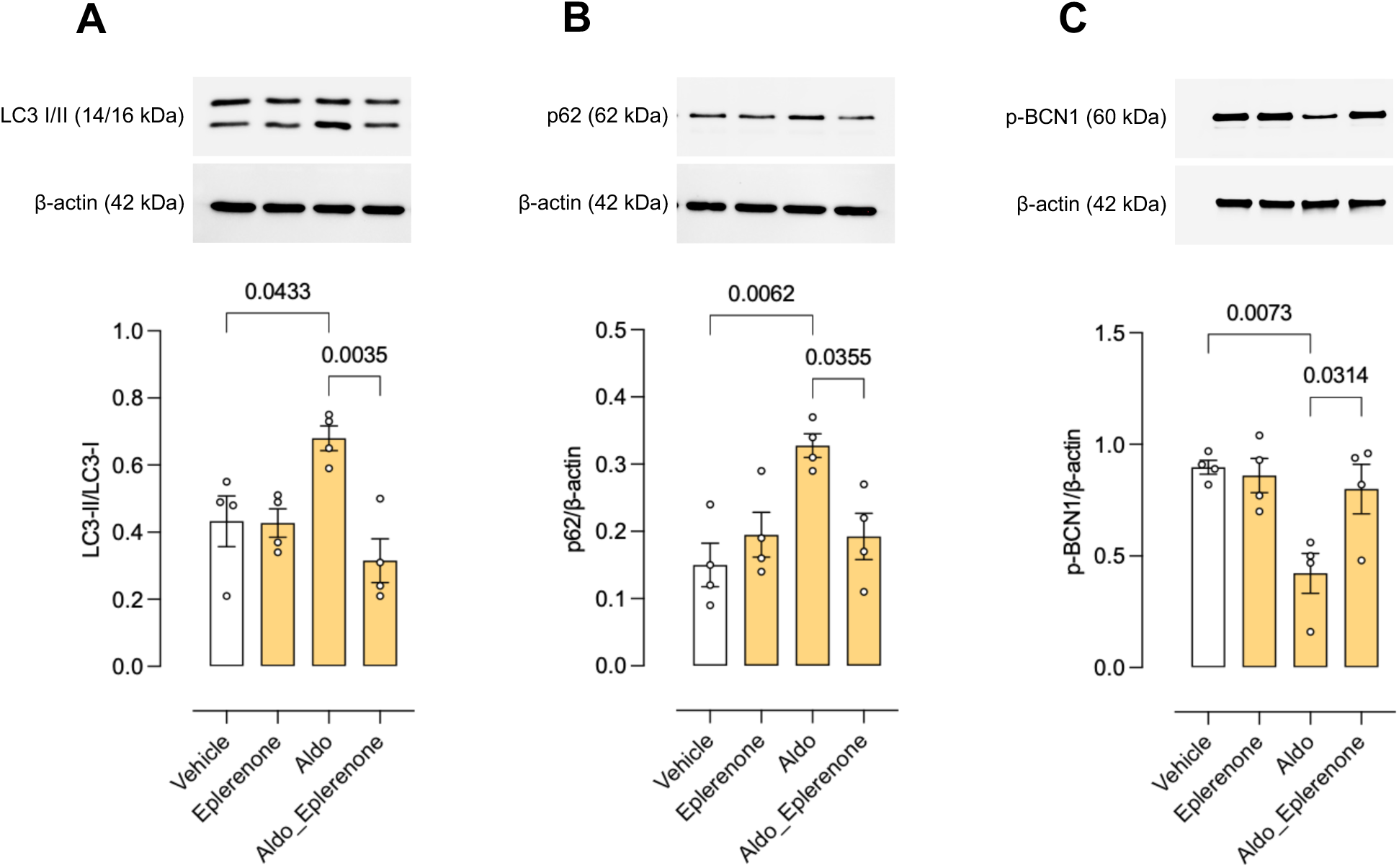
Aldosterone disrupts endothelial autophagy signaling via mineralocorticoid receptor activation. Representative Western blot images showing the expression of LC3 I/II (A), p62 (B), and phosphorylated Beclin-1 (p-BCN1) (C) in mesenteric endothelial cells (MEC) treated with vehicle or aldosterone (0.1 µM, 3h) in the presence or absence of the mineralocorticoid receptor antagonist eplerenone (1 µM, 30 min). Values represent mean ± SEM (n = 4). Statistical analysis was performed using ANOVA.

### Spermidine, an autophagy inducer, protects from aldosterone-induced vascular dysfunction and restores endothelial NO and ROS levels

To determine whether pharmacological induction of autophagy protects against aldosterone-induced vascular dysfunction, mice were treated with spermidine. Autophagy induction significantly improved endothelial function in aldosterone-treated male mice and restored NO production while reducing ROS generation (Figure 3A–C). To verify that spermidine effectively accelerates autophagy, we examined LC3-I/II and p62 expression in MECs exposed to aldosterone. Spermidine normalized both markers of autophagic flux, confirming its ability to restore autophagy signaling disrupted by aldosterone (Figure 3D–E).

**Figure 3.**
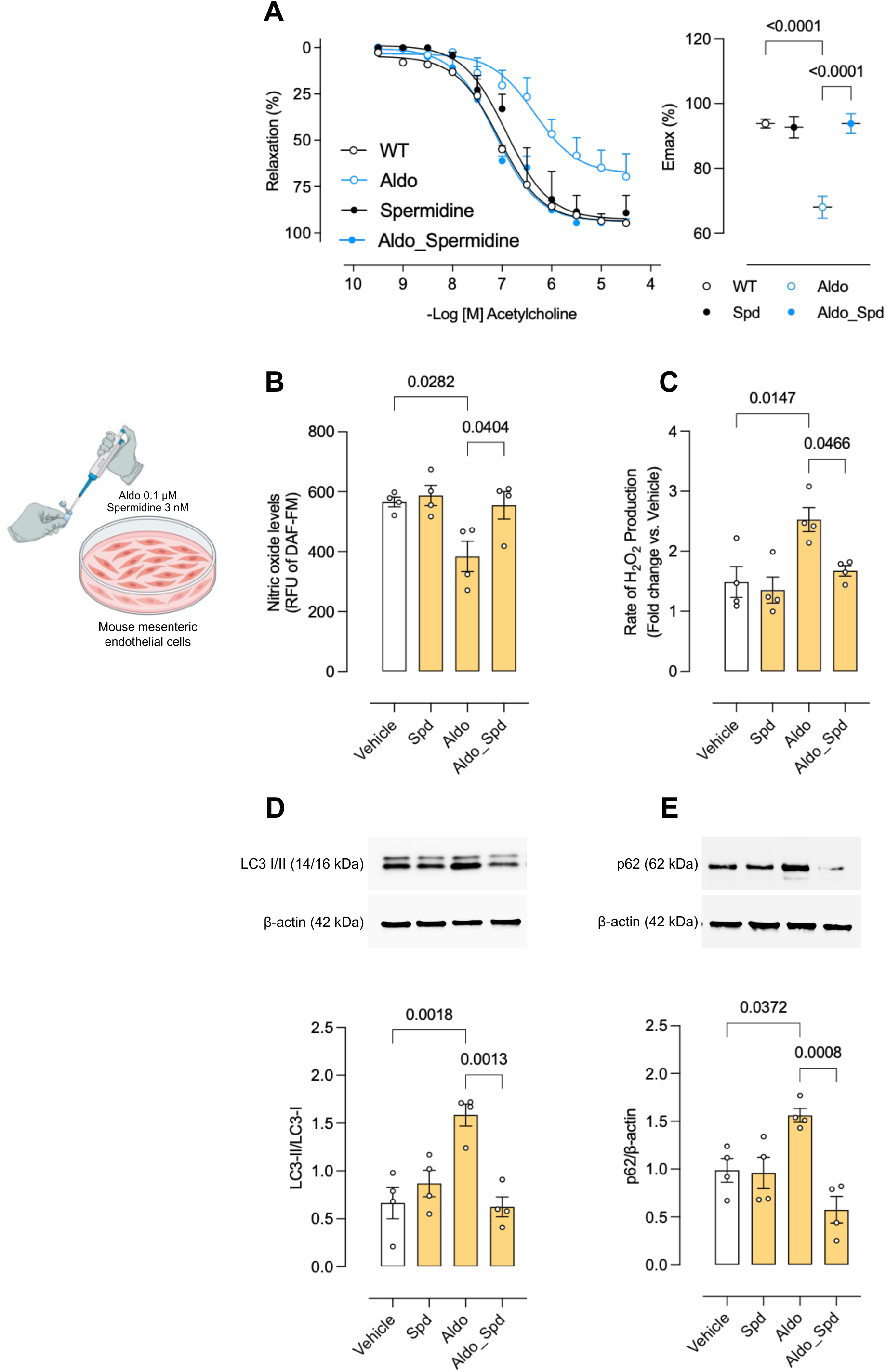
Spermidine restores endothelial function and autophagy signaling impaired by aldosterone. Concentration–response curves to phenylephrine in second-order mesenteric resistance arteries from male wild-type mice treated with vehicle, aldosterone (600 µg/kg/day for 14 days) or spermidine (3 mM in drinking water for 7 days) (A). Nitric oxide production (B) and hydrogen peroxide levels (C) in mesenteric endothelial cells (MEC) treated with vehicle, aldosterone (0.1 µM, 3 h) or spermidine (3 nM, 30 min). Representative Western blot images and densitometric quantification of LC3 I/II (D) and p62 (E) expression in MEC treated with vehicle or aldosterone or spermidine. Values represent mean ± SEM (n = 4–7). Statistical analysis was performed using ANOVA.

### Genetic activation of BCN1-dependent autophagy protects against aldosterone-induced vascular dysfunction and cardiac injury

Next, we utilized BCN1 gain-of-function mice to mechanistically evaluate whether accelerating autophagy confers protection against aldosterone-induced vascular dysfunction and cardiac injury. Male BCN1-overexpressing mice were resistant to aldosterone-induced endothelial dysfunction (Figure 4A). Mesenteric arteries from these mice also demonstrated enhanced autophagic flux, evidenced by increased LC3-I/II levels and preserved p62 expression (Figure 4B–C). Finally, assessment of end-organ damage revealed that aldosterone induced coronary and myocardial fibrosis in control mice, whereas BCN1-overexpressing mice were protected from this cardiac injury (Figure 4D).

**Figure 4.**
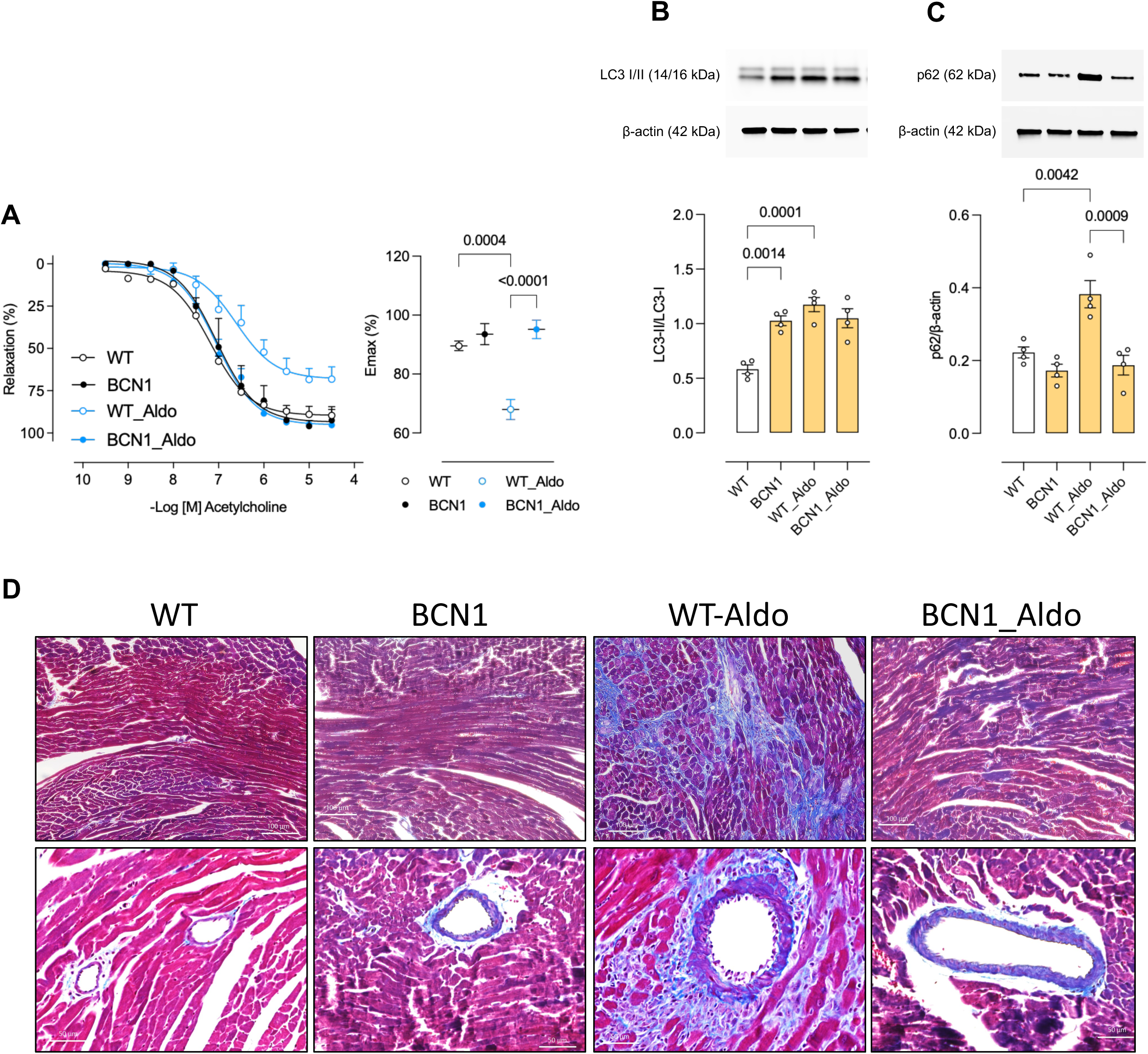
Genetic activation of autophagy protects against aldosterone-induced vascular dysfunction and cardiac injury. Concentration–response curves to acetylcholine in second-order mesenteric resistance arteries (MRA) from male Becn1^F121A^ knock-in mice treated with vehicle or aldosterone (600 µg/kg/day for 14 days) (A). Representative Western blot images and densitometric quantification of LC3 I/II (B) and p62 (C) expression in MRA from Becn1^F121A^ knock-in mice with vehicle or aldosterone. Representative images from hearts of wild type and Becn1^F121A^ knock-in mice treated with vehicle or aldosterone. Masson’s Trichrome was used to stain the fibrotic area (D). Values represent mean ± SEM (n = 4–7). Statistical analysis was performed using ANOVA.

### Pharmacologic BCN1 activation with TB-peptide protects against aldosterone-induced vascular dysfunction and cardiac injury

To provide a pharmacological and translational complement to our genetic model, we used the TB-peptide to activate BCN1 in vivo. TB-peptide treatment improved endothelial function and markedly reduced both coronary and myocardial fibrosis in aldosterone-treated mice (Figure 5A–B). We then confirmed that TB-peptide enhances autophagic flux in MECs. TB-peptide increased LC3-I/II expression without altering p62 levels and restored autophagy in MECs exposed to aldosterone (Figure 5C–D).

**Figure 5.**
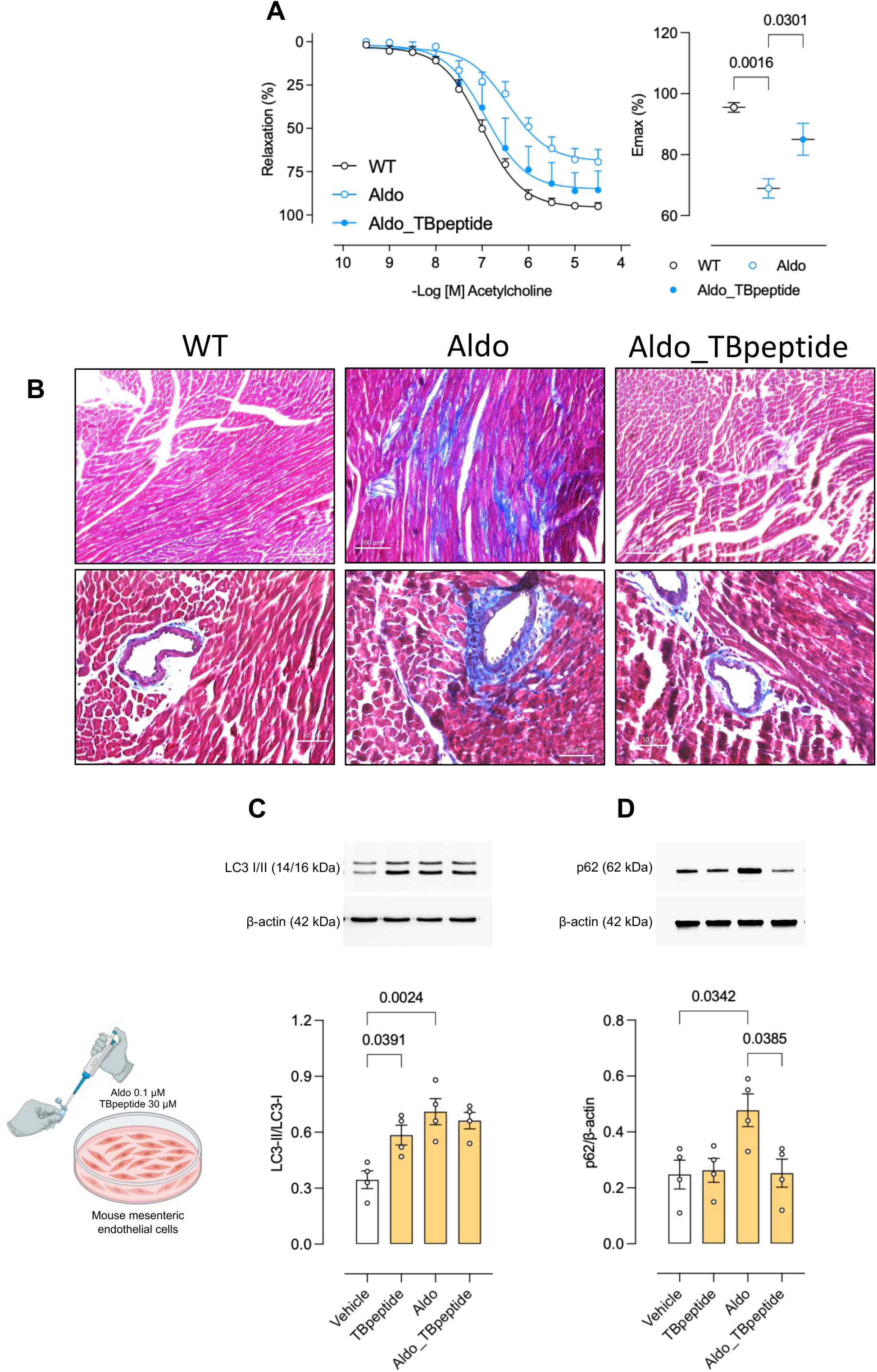
Pharmacological activation of autophagy protects against aldosterone-induced vascular dysfunction and cardiac injury. Concentration–response curves to acetylcholine in second-order mesenteric resistance arteries (MRA) from male wild type mice treated with vehicle or aldosterone (600 µg/kg/day for 14 days) or TB-peptide (16 mg/kg/day for 7 days) (A). Representative images from hearts of male wild type mice treated with vehicle or aldosterone or TB-peptide. Masson’s Trichrome was used to stain the fibrotic area (B). Representative Western blot images and densitometric quantification of LC3 I/II (C) and p62 (D) expression in MEC treated with vehicle or aldosterone (0.1 µM, 3 h) or TB-peptide (30 μM, 30 min). Values represent mean ± SEM (n = 4–7). Statistical analysis was performed using ANOVA.

### BCN1-dependent autophagy preserves endothelial NO/ROS homeostasis and regenerative capacity

Mechanistically, TB-peptide restored NO production in MECs exposed to aldosterone and prevented aldosterone-induced increases in ROS generation (Figure 6A–B). We next assessed endothelial regenerative capacity by evaluating MEC migration. Aldosterone markedly impaired migration in both scratch and transwell assays, whereas TB-peptide treatment significantly rescued migratory ability (Figure 6C–D). Together, these findings indicate that autophagy is critical for maintaining endothelial function and represents a potent strategy to counteract the harmful vascular effects of aldosterone.

**Figure 6.**
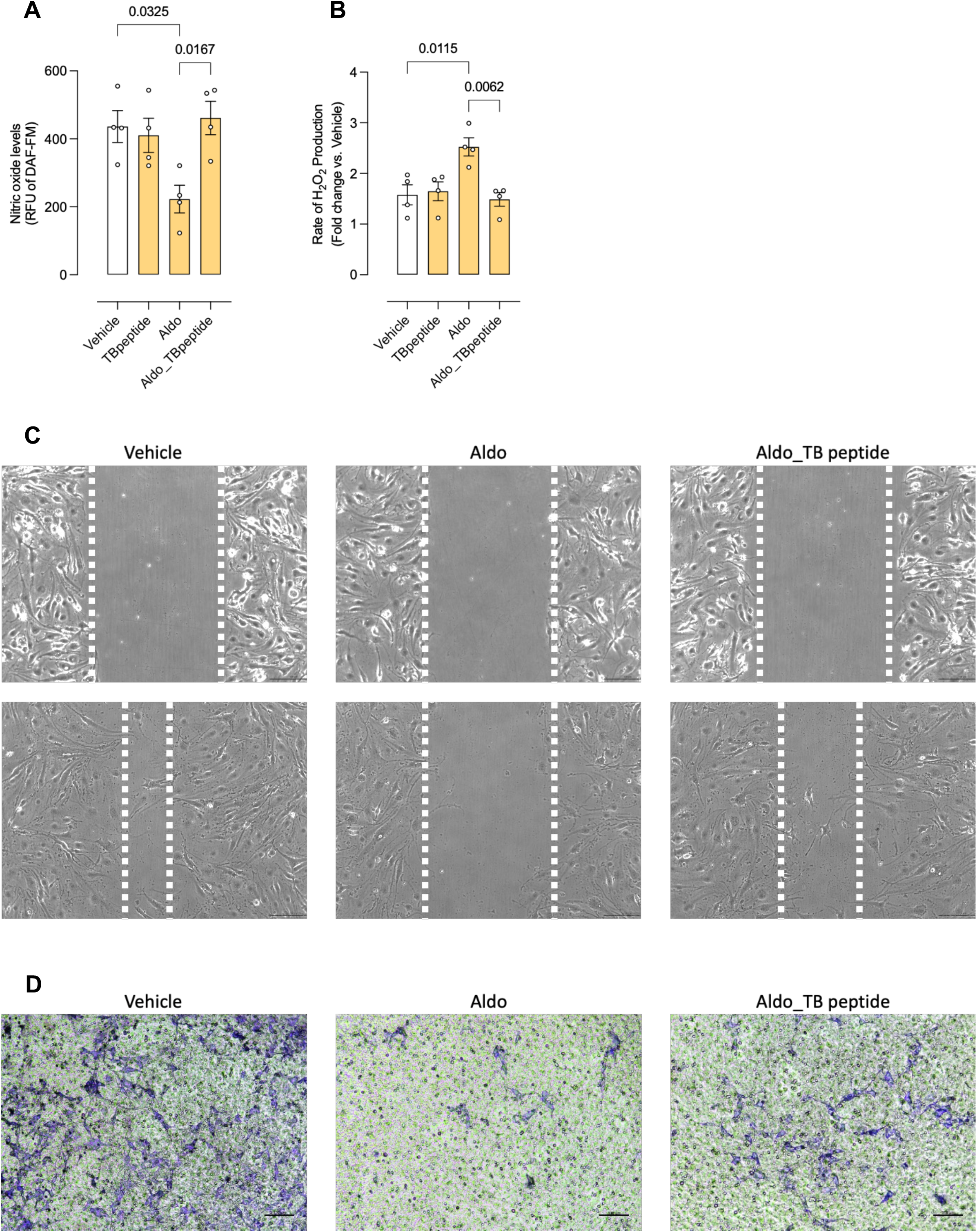
TB-peptide restores endothelial redox balance and migratory capacity impaired by aldosterone. Nitric oxide production (A) and hydrogen peroxide levels (B) in mesenteric endothelial cells (MEC) treated with vehicle, aldosterone (0.1 µM, 3 h) or TB-peptide (30 µM, 30 min). Representative images of MEC migration assessed by scratch wound (C) and transwell (D) assay in MEC treated with vehicle, aldosterone, or aldosterone plus TB-peptide. Values represent mean ± SEM (n = 4). Statistical analysis was performed using ANOVA.

## Discussion

In this study, we demonstrate that impairment of autophagy is a central mechanism contributing to aldosterone-induced endothelial dysfunction, oxidative stress, and cardiovascular injury. Our findings reveal that aldosterone disrupts autophagic flux through an MR-dependent pathway, leading to reductions in NO bioavailability, elevations in ROS production, and diminished endothelial regenerative capacity. Importantly, both pharmacologic and genetic approaches to enhance BCN1-dependent autophagy protected the vasculature from aldosterone-induced injury and prevented downstream cardiac fibrosis. Together, these results identify autophagy activation, specifically through BCN1, as a critical protective mechanism in mineralocorticoid-mediated cardiovascular pathology.

Aldosterone excess is a well-established contributor to vascular dysfunction and hypertension, largely through MR-dependent increases in oxidative stress, inflammation, and impaired endothelial signaling^1–3, 6, 9, 21^. Here, we extend these findings by showing that aldosterone also disrupts autophagic flux in endothelial cells, as evidenced by LC3 accumulation, reduced p62 turnover, and decreased BCN1 expression. MR antagonism fully prevented these alterations, indicating that autophagy impairment is a direct downstream consequence of MR activation. These findings are consistent with recent reports indicating that MR signaling regulates autophagy in metabolic and vascular tissues^6, 22, 23^. Our study provides new evidence that endothelial autophagy is not merely altered by aldosterone but is functionally linked to its detrimental vascular effects.

Restoring autophagy through spermidine treatment significantly improved endothelial function in aldosterone-treated mice and normalized NO and ROS levels in MECs. These findings are consistent with previous work demonstrating that spermidine enhances cardiovascular resilience, improves metabolic homeostasis, and supports endothelial NO signaling^9, 16, 24^. Our results add a new aspect by showing that spermidine can counteract aldosterone-induced vascular dysfunction, highlighting its potential as a therapeutic strategy in conditions characterized by mineralocorticoid excess.

To further dissect the mechanistic contribution of autophagy, we employed BCN1 gain-of-function mice. Enhanced autophagic flux in these animals conferred robust protection against aldosterone-induced endothelial dysfunction and prevented both coronary and myocardial fibrosis. Prior studies have shown that BCN1 overactivation promotes tissue resilience in models of sepsis, metabolic stress, cardiac and PVAT dysfunction, and improves blood pressure^9, 14, 17, 18^. Our findings extend these observations by establishing BCN1-mediated autophagy as a key determinant of vascular health in the context of aldosterone excess, particularly in resistance vessels, which had not previously been examined.

While we previously showed that Beclin1 overexpression partially mitigates aldosterone-induced hypertension9, blood pressure was not evaluated in the present study. Here, our goal was to define vascular-specific mechanisms of protection. The current findings reveal that Beclin1 activation improves endothelial function, restores NO/ROS homeostasis, enhances regenerative capacity, and prevents cardiac fibrosis, indicating that the benefits of autophagy activation extend beyond systemic hemodynamic effects. These results complement our earlier work and highlight autophagy as a fundamental determinant of vascular integrity during mineralocorticoid excess.

Importantly, the use of TB-peptide provided a pharmacological bridge toward translational application. TB-peptide activation of BCN1 restored endothelial function, normalized NO and ROS levels, and reversed aldosterone-induced migration defects. The improvement in endothelial motility is particularly notable, as endothelial regeneration is essential for maintaining vascular integrity following injury. Aldosterone impaired both scratch wound closure and transwell migration, whereas TB-peptide fully rescued these processes, indicating that restoration of autophagy preserves endothelial regenerative capacity and may represent a therapeutic strategy to improve vascular repair. These findings reproduce emerging evidence that autophagy regulates cytoskeletal dynamics, cell motility, and angiogenic responses in endothelial cells^22, 25–27^.

Collectively, our data support a model in which aldosterone-induced MR activation suppresses autophagic flux, thereby promoting oxidative stress, impairing NO bioavailability, and limiting endothelial repair. Restoring autophagy through genetic or pharmacological activation of BCN1 disrupts this pathological signaling pathway, normalizes endothelial homeostasis, and prevents the development of cardiovascular injury. These findings place BCN1-dependent autophagy not only as a biomarker of endothelial health but also as a promising therapeutic target for mineralocorticoid-associated vascular disease.

Several limitations should be acknowledged. First, while our autophagy assays provide evidence of improved autophagic flux, more detailed interrogation using tandem fluorescence reporters or lysosomal blockade could further clarify the step(s) at which aldosterone disrupts autophagy. Second, although we focused on endothelial cells and resistance arteries, aldosterone also affects vascular smooth muscle cells, immune cells, and PVAT^2, 3, 9, 21, 28^. The relative contribution of autophagy in these compartments deserves further investigation. Finally, long-term studies are needed to determine whether sustained BCN1 activation can prevent or reverse structural vascular remodeling beyond what we observed in two weeks of aldosterone exposure.

In conclusion, our study reveals that autophagy impairment is a key mechanism underlying aldosterone-induced vascular dysfunction and cardiac injury. Enhancing BCN1-dependent autophagy, through spermidine or TB-peptide, or via genetic overactivation, restores endothelial NO/ROS balance, preserves vascular regenerative capacity, and prevents end-organ damage. These findings position BCN1-dependent autophagy as a central determinant of endothelial resilience and identify autophagy activation as a promising therapeutic strategy for cardiovascular diseases associated with mineralocorticoid excess and underscore the broader importance of autophagic homeostasis in preserving vascular health.

## Clinical Perspective

- Aldosterone excess contributes to endothelial dysfunction and cardiovascular injury through mineralocorticoid receptor (MR) activation; however, the role of impaired autophagy in this process remains poorly defined. This study was undertaken to determine whether aldosterone disrupts endothelial autophagic flux and whether restoring autophagy could protect vascular and cardiac function.
- We demonstrate that aldosterone impairs autophagy, reduces nitric oxide bioavailability, increases oxidative stress, and induces vascular dysfunction and cardiac fibrosis. Genetic (Beclin1 gain-of-function) and pharmacologic (spermidine and TB-peptide) activation of autophagy restored endothelial homeostasis, improved vascular function, and reduced cardiac injury.
- These findings identify suppression of autophagy as a key mechanism of aldosterone-induced cardiovascular damage and establish Beclin1-dependent autophagy as a therapeutic target. Enhancing autophagy may represent a novel strategy to treat cardiovascular diseases associated with mineralocorticoid excess, including hypertension.

## Data Availability

The datasets generated and/or analyzed during the current study are available from the corresponding author on reasonable request.

## Disclosures

The authors declare no conflicts of interest.

## Acknowledgments

We thank the instrumentation and technical support provided by the University of Pittsburgh and the Department of Pediatrics at UPMC Children’s Hospital.

## Funding

Sao Paulo Research Foundation (FAPES, 2021/08847-9) to RMC; National Heart, Lung, and Blood Institute (NHLBI, R01-HL169202) to TBN. Startup funds from University of Pittsburgh to TBN; Startup funds from University of South Alabama to TBN.

## Notes

### Competing Interest Statement

The authors have declared no competing interest.

## References

1. Buffolo F, Tetti M, Mulatero P, Monticone S. Aldosterone as a Mediator of Cardiovascular Damage. Hypertension. 2022;79(9):1899–911.

2. Bruder-Nascimento T, Ferreira NS, Zanotto CZ, Ramalho F, Pequeno IO, Olivon VC, et al. NLRP3 Inflammasome Mediates Aldosterone-Induced Vascular Damage. Circulation. 2016;134(23):1866–80.

3. Costa RM, Cerqueira DM, Bruder-Nascimento A, Alves JV, Awata WMC, Singh S, et al. Role of the CCL5 and Its Receptor, CCR5, in the Genesis of Aldosterone-Induced Hypertension, Vascular Dysfunction, and End-Organ Damage. Hypertension. 2024;81(4):776–86.

4. Jaffe IZ, Jaisser F. Endothelial cell mineralocorticoid receptors: turning cardiovascular risk factors into cardiovascular dysfunction. Hypertension. 2014;63(5):915–7.

5. Bauersachs J, Lother A. Mineralocorticoid receptor activation and antagonism in cardiovascular disease: cellular and molecular mechanisms. Kidney Int Suppl (2011). 2022;12(1):19–26.

6. Wang M, Wang C, Zhao M, Feng Z, Li Y, Lan L, et al. Mineralocorticoid receptor regulates vascular damage by inducing autophagy in mice with obesity and hypertension. Nutr Metab Cardiovasc Dis. 2025;35(11):104191.

7. Nakahira K, Cloonan SM, Mizumura K, Choi AM, Ryter SW. Autophagy: a crucial moderator of redox balance, inflammation, and apoptosis in lung disease. Antioxid Redox Signal. 2014;20(3):474–94.

8. Li A, Gao M, Liu B, Qin Y, Chen L, Liu H, et al. Mitochondrial autophagy: molecular mechanisms and implications for cardiovascular disease. Cell Death Dis. 2022;13(5):444.

9. Costa RM, Bruder-Nascimento A, Alves JV, Awata WMC, Singh S, Rodrigues D, et al. Beclin-1-dependent autophagy protects perivascular adipose tissue function from hyperaldosteronism effects. Am J Physiol Heart Circ Physiol. 2025;328(6):H1253–H66.

10. Tian Z, Ning H, Wang X, Wang Y, Han T, Sun C. Endothelial Autophagy Promotes Atheroprotective Communication Between Endothelial and Smooth Muscle Cells via Exosome-Mediated Delivery of miR-204-5p. Arterioscler Thromb Vasc Biol. 2024;44(8):1813–32.

11. McCarthy CG, Wenceslau CF, Calmasini FB, Klee NS, Brands MW, Joe B, et al. Reconstitution of autophagy ameliorates vascular function and arterial stiffening in spontaneously hypertensive rats. Am J Physiol Heart Circ Physiol. 2019;317(5):H1013–H27.

12. Kim KA, Shin D, Kim JH, Shin YJ, Rajanikant GK, Majid A, et al. Role of Autophagy in Endothelial Damage and Blood-Brain Barrier Disruption in Ischemic Stroke. Stroke. 2018;49(6):1571–9.

13. Aman Y, Schmauck-Medina T, Hansen M, Morimoto RI, Simon AK, Bjedov I, et al. Autophagy in healthy aging and disease. Nat Aging. 2021;1(8):634–50.

14. Son Y, Cho YK, Saha A, Kwon HJ, Park JH, Kim M, et al. Adipocyte-specific Beclin1 deletion impairs lipolysis and mitochondrial integrity in adipose tissue. Mol Metab. 2020;39:101005.

15. Singh S, Bruder A, Costa RM, Alves JV, Bharathi S, Goetzman ES, et al. Vascular Contractility Relies on Integrity of Progranulin Pathway: Insights Into Mitochondrial Function. J Am Heart Assoc. 2025:e037640.

16. Eisenberg T, Abdellatif M, Schroeder S, Primessnig U, Stekovic S, Pendl T, et al. Cardioprotection and lifespan extension by the natural polyamine spermidine. Nat Med. 2016;22(12):1428–38.

17. Kim M, Nikouee A, Zou R, Ren D, He Z, Li J, et al. Age-Independent Cardiac Protection by Pharmacological Activation of Beclin-1 During Endotoxemia and Its Association With Energy Metabolic Reprograming in Myocardium-A Targeted Metabolomics Study. J Am Heart Assoc. 2022;11(14):e025310.

18. Sun Y, Yao X, Zhang QJ, Zhu M, Liu ZP, Ci B, et al. Beclin-1-Dependent Autophagy Protects the Heart During Sepsis. Circulation. 2018;138(20):2247–62.

19. Bruder-Nascimento A, Awata WMC, Alves JV, Singh S, Costa RM, Bruder-Nascimento T. Progranulin Maintains Blood Pressure and Vascular Tone Dependent on EphrinA2 and Sortilin1 Receptors and Endothelial Nitric Oxide Synthase Activation. J Am Heart Assoc. 2023;12(16).

20. Alves JV, da Costa RM, Awata WMC, Bruder-Nascimento A, Singh S, Tostes RC, et al. NADPH oxidase 4-derived hydrogen peroxide counterbalances testosterone-induced endothelial dysfunction and migration. Am J Physiol-Endoc M. 2024;327(1):E1–E12.

21. Bruder-Nascimento T, Callera GE, Montezano AC, Belin de Chantemele EJ, Tostes RC, Touyz RM. Atorvastatin inhibits pro-inflammatory actions of aldosterone in vascular smooth muscle cells by reducing oxidative stress. Life Sci. 2019;221:29–34.

22. Armani A, Cinti F, Marzolla V, Morgan J, Cranston GA, Antelmi A, et al. Mineralocorticoid receptor antagonism induces browning of white adipose tissue through impairment of autophagy and prevents adipocyte dysfunction in high-fat-diet-fed mice. FASEB J. 2014;28(8):3745–57.

23. Habibi J, Chen D, Hulse JL, Whaley-Connell A, Sowers JR, Jia G. Targeting mineralocorticoid receptors in diet-induced hepatic steatosis and insulin resistance. Am J Physiol Regul Integr Comp Physiol. 2022;322(3):R253–R62.

24. Stojanovic I, Jelenkovic A, Stevanovic I, Pavlovic D, Bjelakovic G, Jevtovic-Stoimenov T. Spermidine influence on the nitric oxide synthase and arginase activity relationship during experimentally induced seizures. J Basic Clin Physiol Pharmacol. 2010;21(2):169–85.

25. Shadab M, Millar MW, Slavin SA, Leonard A, Fazal F, Rahman A. Autophagy protein ATG7 is a critical regulator of endothelial cell inflammation and permeability. Sci Rep. 2020;10(1):13708.

26. Torisu T, Torisu K, Lee IH, Liu J, Malide D, Combs CA, et al. Autophagy regulates endothelial cell processing, maturation and secretion of von Willebrand factor. Nat Med. 2013;19(10):1281–7.

27. Schaaf MB, Houbaert D, Mece O, Agostinis P. Autophagy in endothelial cells and tumor angiogenesis. Cell Death Differ. 2019;26(4):665–79.

28. Kasal DA, Barhoumi T, Li MW, Yamamoto N, Zdanovich E, Rehman A, et al. T regulatory lymphocytes prevent aldosterone-induced vascular injury. Hypertension. 2012;59(2):324–30.

